# Lopinavir-ritonavir is not an effective inhibitor of the main protease activity of SARS-CoV-2 *in vitro*

**DOI:** 10.1101/2020.09.16.299800

**Authors:** Minsu Jang, Yea-In Park, Rackhyun Park, Yeo-Eun Cha, Sim Namkoong, Jin I. Lee, Junsoo Park

## Abstract

COVID-19 has caused over 900,000 deaths worldwide as of September 2020, and effective medicines are urgently needed. Lopinavir was identified as an inhibitor of the HIV protease, and a lopinavir-ritonavir combination therapy was reported to be beneficial for the treatment of SARS and MERS. However, recent clinical tests could not prove that lopinavir-ritonavir therapy was an effective treatment for COVID-19. In this report, we examined the effect of lopinavir and ritonavir to the activity of the purified main protease (Mpro) protein of SARS-CoV-2, the causative virus of COVID-19. Unexpectedly, lopinavir and ritonavir did not inhibit Mpro activity. These results will aid the drug candidate selection for ongoing and future COVID-19 clinical trials.

## 1. Introduction

Lopinavir is an approved drug for treatment against human immunodeficiency virus-1 (HIV-1) and works by inhibiting the HIV-1 protease [1]. The approved drug ritonavir increases the plasma level of lopinavir dramatically by inhibiting cytochrome p450 [1]. When severe acute respiratory syndrome (SARS) emerged in the 2000s, lopinavir was identified as a candidate drug to treat SARS-coronavirus (SARS-CoV) [2]. Lopinavir showed an inhibitory effect against SARS-CoV in *in vitro* studies, and clinical tests with the lopinavir-ritonavir combination showed a reduced risk of SARS-CoV infection [3, 4]. Lopinavir-ritonavir treatment was also beneficial to treat Middle East respiratory syndrome coronavirus (MERS-CoV) [5, 6]. SARS-CoV-2, the virus responsible for the current COVID-19 outbreak, is closely related to and shows high sequence homology with SARS-CoV and, to a slightly lesser extent, MERS-CoV coronaviruses [7]. For this reason, several clinical tests for COVID-19 were performed with lopinavir-ritonavir, however the clinical results have not proved that lopinavir-ritonavir treatment was beneficial to treat COVID-19 [8, 9].

Lopinavir is known to be an inhibitor of the SARS-CoV main protease (Mpro, also known as 3CL protease), and recent bioinformatics studies showed that lopinavir may interact with Mpro of SARS-CoV-2 [10, 11]. For this reason, lopinavir was assumed to inhibit Mpro of SARS-CoV-2, however no evidence of this has been presented. In light of the recent COVID-19 clinical tests with lopinavir-ritonavir treatment, we attempted to evaluate the inhibitory activity of lopinavir for Mpro of SARS-CoV-2. Using an *in vitro* protease assay with the purified Mpro protein demonstrated that lopinavir as well as ritonavir, we found that two drugs did not have any inhibitory activity against Mpro of SARS-CoV-2. This study highlights differences between the highly related SARS, MERS and COVID-19-causing coronaviruses, and will be helpful in the selection of drug candidates for current and future COVID-19 clinical tests.

## 2. Material and methods

### 2.1. Reagents

Lopinavir (SML1222, purity ≥ 98 %), ritonavir (SML0491, purity ≥ 98 %), and (−)-Epigallocatechin gallate (EGCG) (E4134, purity ≥ 95 %) were purchased from Sigma-Aldrich (Saint Louis, MO).

### 2.2. Protease assay for SARS-COV-2 Mpro

Preparation of SARS-CoV-2 Mpro was described previously [12]. Briefly, the nucleotide sequence of SARS-CoV-Mpro was chemically synthesized and His-tagged Mpro was expressed in BL21(DE3) cells, and soluble Mpro proteins were purified with an Ni-NTA resin (Thermo-Fisher Scientific, Rockford, IL) according to the manufacturer’s protocol. A FRET-based protease assay was used to measure Mpro activity [12, 13]. Dabcyl-KTSAVLQSGFRKME-Edans was chemically synthesized (Anygen, Gwangju, South Korea) and used for SARS-COV-2 Mpro substrate. The protease activity was performed using Mpro and FRET peptide in the reaction buffer (20 mM Tris-HCl (pH 7.5), 200 mM NaCl, 5 mM EDTA, 5 mM DTT, 1 % DMSO) for 5 h. For the inhibition assay, the purified Mpro protein was incubated with the chemicals for 1 h before the addition of substrate. Relative protease activity was calculated as the difference between the protease activity with Mpro and the activity without Mpro at the indicated time.

## 3. Results

Lopinavir/ritonavir treatment is currently being studied in several clinical trials for COVID-19, and, in light of these studies, we attempted to measure the effect of lopinavir/ritonavir on SARS-CoV-2. First, we set up the Mpro protease assay system with purified Mpro protein from bacteria (Figure 1B and 1C). Next, Mpro protein was mixed with various concentration of lopinavir, and the protease activity of Mpro was measured. Although high concentrations of lopinavir up to 100 μg/ml were used, we did not observe any significant decrease of Mpro activity (Figure 1D). Because lopinavir and ritonavir combination therapy is commonly used for the clinical trials, we also examined the inhibitory activity of lopinavir and ritonavir combinations. However, the lopinavir-ritonavir combination treatment also did not show any significant decrease of Mpro activity (Figure 1E). As a positive control for MPro activity, we used the compound EGCG [12] which showed significant inhibition of Mpro activity (Figure 1F). These results indicate that lopinavir and ritonavir did not have an inhibitory effect on Mpro activity of SARS-CoV-2.

**Figure 1.**
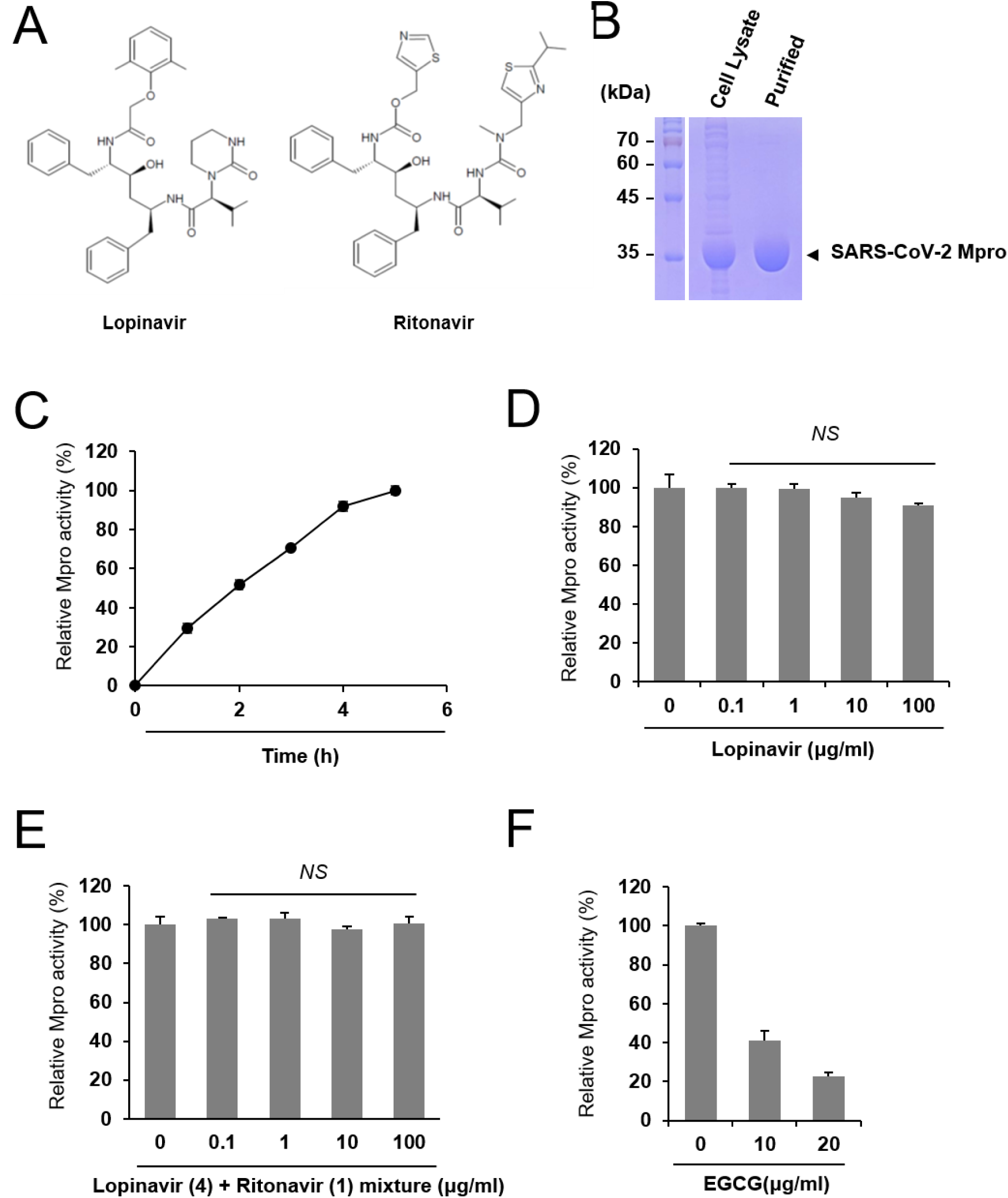
Lopinavir is ineffective to inhibit SARS-CoV-2 Mpro. (A) Chemical structure of lopinavir and ritonavir. (B) Purification of SARS-CoV-2 Mpro. Cell lysate and purified Mpro protein were subject to SDS-PAGE. (C) Measurement of the relative Mpro activity over time. Activity is measured relative to maximal activity at 5 hr. (D) Lopinavir did not inhibit Mpro. Indicated concentrations of lopinavir were incubated with Mpro and the relative protease activity was determined. The Mpro activity was measured in quadruple and the mean and standard deviation are shown. NS, not significant. (E) Mixtures of lopinavir and ritonavir at a weight ratio of 4:1 were incubated with Mpro, and the relative protease activity was determined. (F) EGCG was incubated with Mpro, and the protease activity was examined as a positive control.

## 4. Discussion

Lopinavir-ritonavir is a candidate medicine for COVID-19, however clinical trials with lopinavir-ritonavir did not prove that these medicines are beneficial for COVID-19 patients [8]. Here, we examined the inhibitory effect of lopinavir-ritonavir on SARS-CoV-2 Mpro protease activity and found that lopinavir-ritonavir is not effective to inhibit SARS-CoV-2 Mpro activity. Lopinavir was reported to inhibit SARS-CoV Mpro with an IC50 of 25∼50 μM [2]. However, lopinavir nor combinations of lopinavir and ritonavir were not effective to inhibit SARS-CoV- 2 Mpro at any of the concentrations we tested. These results may explain the difference in drug response on SARS and COVID-19 patients in the clinical trials.

Since lopinavir responds differently to SARS-CoV Mpro and SARS-CoV-2 Mpro, we wondered if structural differences between the two Mpro proteins could account for the disparity. Previous computational analysis studies listed the potential amino acid residues responsible for the interaction between Mpro and lopinavir [10, 14-16]. Because SARS-CoV- 2 Mpro shares 96.08% identity with SARS-CoV Mpro, most of the listed amino acids are identical [12]. Among the potential amino acids responsible for the interaction with lopinavir, we found just one amino acid difference. Serine residue 46 of SARS-CoV-2 Mpro is one of the candidate amino acids for the interaction with lopinavir [10], however both SARS-CoV and MERS-CoV bear an alanine residue at that site in their Mpro protein sequences. Although the sequence differences are not substantial, these trivial sequence differences may contribute to the disparity in Mpro response to lopinavir. Further structural study will be required to reveal the significance of these amino acid changes.

Because a remedy for COVID-19 is very urgent, many clinical trials for COVID-19 are under way. Conducting clinical trials consumes tremendous time, effort and resource, target drug information including *in vitro* data such as those presented here will be very important for the success of the clinical trials. In this report, we demonstrated that lopinavir-ritonavir is not effective to inhibit SARS-CoV-2 Mpro, and this information will be useful to choose the right drug for SARS-CoV-2 clinical trials.

## Acknowledgements

This study was supported by a National Research Foundation of Korea (NRF) grant funded by the Korean government (2019R1A2C1006511).

## Abbreviations

COVID-19: the 2019 novel coronavirus disease
SARS-CoV-2: severe acute respiratory syndrome coronavirus 2
Mpro: main protease,
EGCG: (-)-epigallocatechin-3-gallate

